# *Aspergillus niger* citrate exporter revealed by comparison of two alternative citrate producing conditions

**DOI:** 10.1101/259051

**Authors:** Dorett I Odoni, Thanaporn Laothanachareon, Marta Vazquez-Vilar, Merlijn P van Gaal, Tom Schonewille, Lyon Bruinsma, Vitor AP Martins dos Santos, Juan Antonio Tamayo-Ramos, Maria Suarez-Diez, Peter J Schaap

**Author notes:** Corresponding authors: Dorett I Odoni (DIO);, Peter J Schaap; (PJS). These authors contributed equally to this work (Dorett I Odoni, Thanaporn Laothanachareon, Marta Vazquez-Vilar).

## Abstract

Currently, there is no consensus regarding the mechanism underlying *Aspergillus niger* citrate biosynthesis and secretion, although it is amongst the most studied biotechnological production processes. Carbon excess relative to various other medium constituents is key, but the complex interplay between the limiting factors required for extracellular citrate accumulation remains elusive. It is thought that one of the industrial bottlenecks for citrate production is citrate export, however, no *A. niger* citrate exporter has yet been identified. Here, we show that the phenotype of increased extracellular citrate accumulation can have fundamentally different underlying mechanisms, depending on how this response is triggered, and that combining gene expression analyses of the different conditions can lead to the compilation of a shortlist of the most promising citrate exporter candidates. Specifically, we found that varying the amount and type of supplement of an arginine auxotrophic *A. niger* strain shows down-regulation of citrate metabolising enzymes in the condition in which more citrate is accumulated extracellularly. This contrasts with the transcriptional adaptations triggered by iron limitation, which also induces increased *A. niger* citrate production. By combining data obtained from these two manners of inducing comparatively high extracellular citrate accumulation, we were able to compile a shortlist of the most likely citrate transporter candidates. Two of the most promising candidates were tested in the yeast *Saccharomyces cerevisiae*, one of which showed the ability to secrete citrate. Deletion of the endogenous *A. niger* gene encoding the corresponding transporter abolished the ability of this fungus to secrete citrate. Instead, under conditions that usually favour *A. niger* citrate production, we found increased accumulation of extracellular oxalate. Our findings provide steps in untangling the complex interplay of different mechanisms underlying *A. niger* citrate accumulation, and we identify, for the first time, a fungal citrate exporter, offering a valuable tool for improvement of *A. niger* as biotechnological cell-factory for organic acid production.

**Author Summary:** Citrate is widely applied as acidifier, flavouring and chelating agent. Industrial citrate production currently relies on the filamentous fungus *Aspergillus niger*. Although the industrial production process using *A*. niger has vastly improved since initiated almost 100 years ago, citrate export remains a bottleneck. Here, we studied the gene expression pattern of *A. niger* under various citrate producing conditions. Using these expression patterns and different computational approaches, we compiled a shortlist of putative citrate exporter candidates. In this way, we were able to identify a gene encoding a transporter protein capable of citrate export. We show that the yeast *Saccharomyces cerevisiae*, normally a citrate non-producer, secretes detectable amounts of citrate when harbouring this gene. In addition, we verify the biological function of this gene in *A. niger* itself, as removing this gene resulted in a citrate non-producing phenotype, which is atypical for this fungus. This finding is particularly exciting, as it is the first identification of a eukaryotic citrate exporter. With this, we not only provide a tool for improvement of industrial citrate production, but knowledge of this gene should help develop new methods for improvement of *A. niger* as biotechnological cell-factory for the production of other organic acids.

## Introduction

Large scale *Aspergillus niger* citrate production is a notorious example of a production process that requires a unique combination of unusual nutrient and environmental conditions [1]. Carbon excess relative to iron, zinc, copper, manganese, phosphorus, magnesium, potassium and nitrogen reportedly all leads to an increase in *A. niger* citrate production [2, 3]. The result of this is that, although *A. niger* citrate production has been subject to study since Curries fundamental breakthroughs regarding *A. niger* citrate fermentation > 100 years ago [4], it is still not fully understood.

One problem might be that there are multiple factors at play, and studies therefore contradict each other depending on which aspect or time point of citrate accumulation was investigated [1]. Citrate secretion has long been accepted to be a response to unfavourable intracellular conditions that lead to excess fluxes through the TCA cycle, and ultimately the undesired (from the perspective of the fungus) accumulation of citrate, *i.e*. overflow metabolism [5]. Nevertheless, the viewpoint that citrate is solely an overflow metabolite is changing, and *A. niger* citrate secretion might also be regarded as a response to environmental conditions, such as competition [6], or low iron bioavailability [7]. While large scale production of citrate can probably be attributed to a combination of different mechanisms, increased understanding of all the various aspects that lead to increased citrate production can provide tools to further control and modulate citrate production.

Another important element of metabolic control can be found at transporter level [1], and overexpressing or introducing specific transporters for the product of interest can provide additional tools to overcome product limitation [8]. Dynamic models of metabolism have highlighted the citrate exporter as one of the proteins whose overexpression could lead to increased citrate production rates [9]. However, even though the citrate transport system in *A. niger* has been described [10], no citrate exporter has yet been identified. Knowledge and characterisation of the *A. niger* citrate exporter would thus open new venues to further control and increase product formation.

Here, we explore how changes in media composition induce changes in citrate production, and how combining multiple gene expression datasets can narrow down the list of putative citrate exporter candidates. For this, we worked with a well-defined *A. niger argB* knock-out mutant. The Δ*argB* mutation induces an arginine auxothropy that can be overcome by media supplementation with either arginine or citrulline [11]. We studied the impact of supplement type and amount on citrate production and performed comparative transcriptome analysis to pinpoint the metabolic adaptations associated with the higher citrate producing condition. We compared the results of this study with previous data regarding changes in citrate production triggered by low iron availability [7]. Differences and similarities in the underlying transcriptomic landscape can help discern universal changes upon induced citrate production, or more specific changes given the condition, and served as the starting point for the identification of putative citrate transporter candidates. We validated our approach by testing two of the most promising citrate exporter candidates in *Saccharomyces cerevisiae*, and show that one of these yeast transformant strains indeed accumulated a measurable amount of extracellular citrate. In a next step, we show that a knock-out of the gene encoding the corresponding transporter abolished citrate secretion in *A. niger*.

## Results and discussion

### Experimental setup for selection of candidate citrate exporters

To study the mechanisms underlying extracellular citrate accumulation, and use these insights to pinpoint putative citrate exporter candidates, we compared gene expression data from two different experimental setups that can induce changes in *A. niger* citrate secretion; iron limitation [7] (referred to as “iron experiment” and indicated with +Fe/-Fe), and citrulline supplementation of an arginine auxotrophic mutant (referred to as “supplement experiment” and indicated with _a/_c) (Fig 1A).

**Fig 1.**
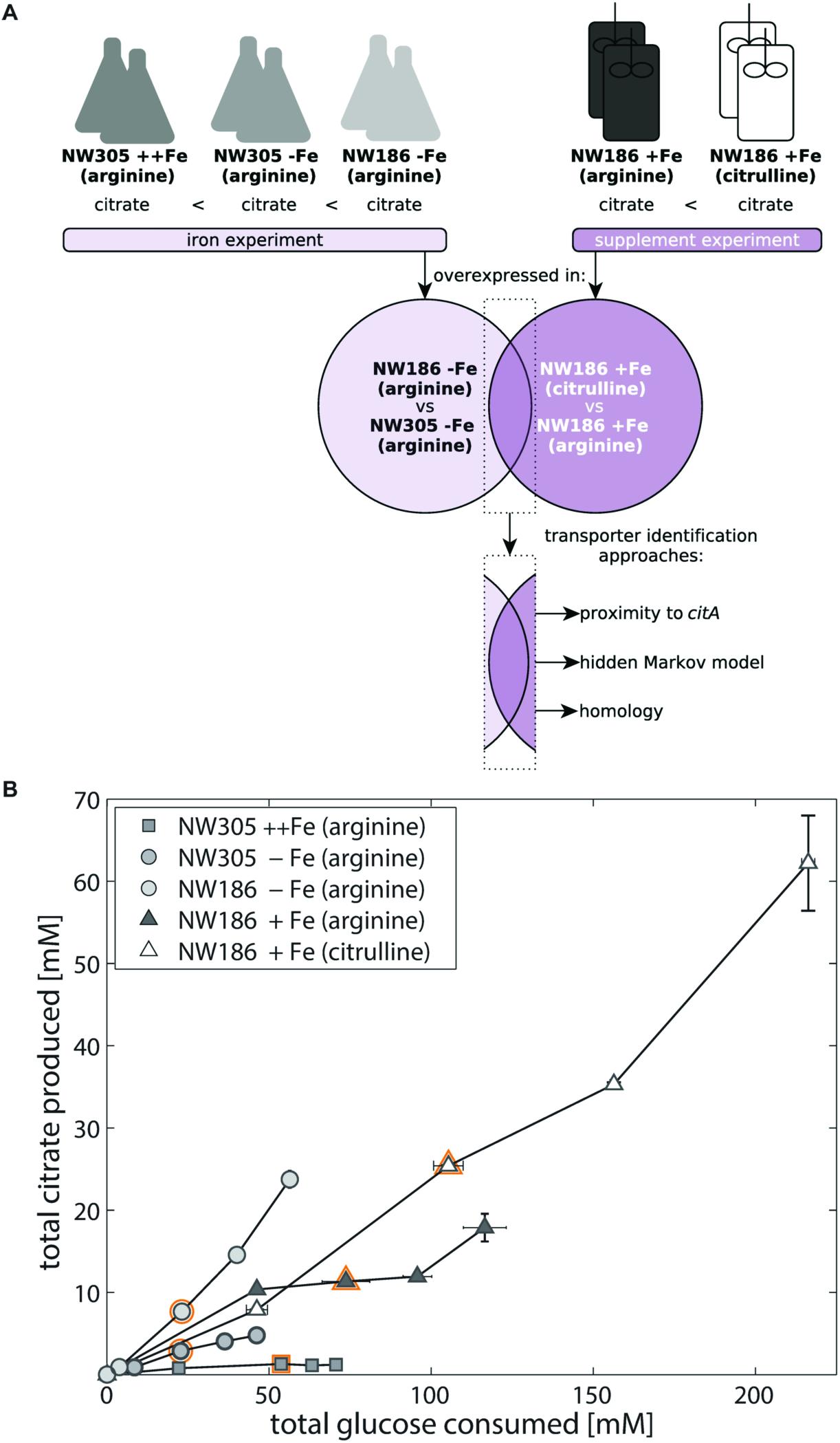
Experimental setup and citrate exporter identification workflow. A) In the iron experiment, performed in shake flasks, extracellular citrate accumulation was modulated by reducing iron availability in the culture medium [7]. In the supplement experiment, performed in fermentors, extracellular citrate accumulation was modulated by substituting the arginine supplement with excess citrulline. Lists of differentially expressed genes in both experimental setups were compared to obtain a shortlist of the most promising citrate exporter candidates. The list of citrate exporter candidates was further narrowed down following the indicated bioinformatic approaches. B) Total citrate production per glucose consumption of NW186 and NW305 grown in shake flasks (iron experiment) or fermentors (supplement experiment), without addition of iron (circles), or varying amounts of Fe(II)SO_4_ (triangles = 1 g·L^-1^, squares = 10 g·L^-1^), and supplemented with either arginine (filled symbols) or excess citrulline (empty symbols). Sample points for RNA extraction (t = 48 h) are marked in orange. Measurement points were taken once every 24 h and show the average of two biological replicates.

In these experiments, two *A. niger* N402 derivative strains (NW186 and NW305) were used. Both NW186 and NW305 have a mutation in glucose oxidase (*goxC17*), making these strains gluconate non-producers. In addition, NW186 has a mutation in oxaloacetate acetylhydrolase (*prtF28*), making this strain an oxalate non-producer [12]. As a result, NW305 is able to produce both oxalate and citrate, whereas NW186 can only produce citrate. Both strains are in addition ornithine transcarbamylase (*argB*) knock-out mutants, making them arginine auxotrophic [11]. Supplementation of the medium with either arginine or citrulline can restore growth of the *ΔargB* mutants, but we found that addition of excess (5 mM) citrulline (NW186 +Fe_c) increases total citrate production when compared to addition of (1.1 mM; “standard” condition) arginine (NW186 +Fe_a). Thus, as can be seen Fig 1B, we can induce an increase in relative citrate production by either limiting iron availability (iron experiment), or substituting the arginine supplement with excess citrulline (supplement experiment).

Citrate production yields of the considered conditions are shown in Table 1. Total citrate yield is almost doubled in NW186 +Fe_c compared to NW186 +Fe_a (Table 1). As expected, addition of excess citrulline does not only increase the citrate productivity (g·L^-1^·h^-1^, Fig 1B), but also has an overall stimulating effect on metabolism; it leads to increased glucose consumption (Fig 1B), doubled final biomass (Table 1), and increased CO_2_ production (337.79 ± 15.86 mM in NW186 +Fe_c, and 265.51 ± 9.28 mM in NW186 +Fe_a). On the other hand, not adding iron to the culture medium increases the citrate per glucose production rate (Fig 1B), but total biomass production remains limited (Table 1), and reduced glucose consumption is observed (Fig 1B). These differences between both experiments suggest that there are two quite different mechanisms at play, although both lead to an increase in extracellular citrate accumulation relative to their respective control conditions.

**Table 1.** Final biomass and citrate yields of *A. niger* NW305 and NW186 grown with varying iron concentrations or supplements in the medium.

| Experimental setup | Strain | Fe | Suppl<sup>2</sup> | Final biomass [g] | Citrate yield |  |
| --- | --- | --- | --- | --- | --- | --- |
|  |  |  |  |  | [g/g biomass] | [g/g substrate] |
| shake flask | NW305<sup>1</sup> | ++ | a | 0.35 ± 0.02 | 0.67 ± 0.07 | 0.02 ± 0.002 |
| shake flask | NW305<sup>1</sup> | - | a | 0.19 ± 0.02 | 4.92 ± 0.35 | 0.11 ± 0.002 |
| shake flask | NW186<sup>1</sup> | - | a | 0.22 ± 1e<sup>-03</sup> | 21.88 ± 1.51 | 0.46 ± 0.02 |
| fermentor | NW186 | + | a | 1.87 ± 0.04 | 1.80 ± 0.14 | 0.17 ± 0.03 |
| fermentor | NW186 | + | c | 3.96 ± 0.09 | 2.98 ± 0.35 | 0.31 ± 0.03 |

<sup>1</sup>data from [7], <sup>2</sup>Suppl: supplements; a: arginine; c: citrulline

### General RNA seq analysis

Transcriptomic data corresponding to the iron experiment have been published in [7]. Here, we performed transcriptomics analysis on the supplement experiment, *i.e*. NW186 grown with Fe(II)SO_4_ added to the medium, and supplemented with either 1.1 mM arginine or 5 mM citrulline. RNA for RNA sequencing (RNA seq) was extracted after 48 h of growth. The annotated genome of *A. niger* ATCC 1015 [13] was used as reference to map the RNA seq reads (Table 2). Genes with count per million (CPM) ≥ 1 were considered to be expressed (S1 Dataset). The supplement change from arginine to excess citrulline induces major transcriptional adaptations, and over 20% of the annotated genes are differentially expressed (Table 2, S2 Dataset).

**Table 2.**
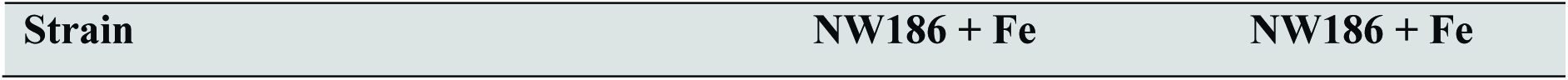
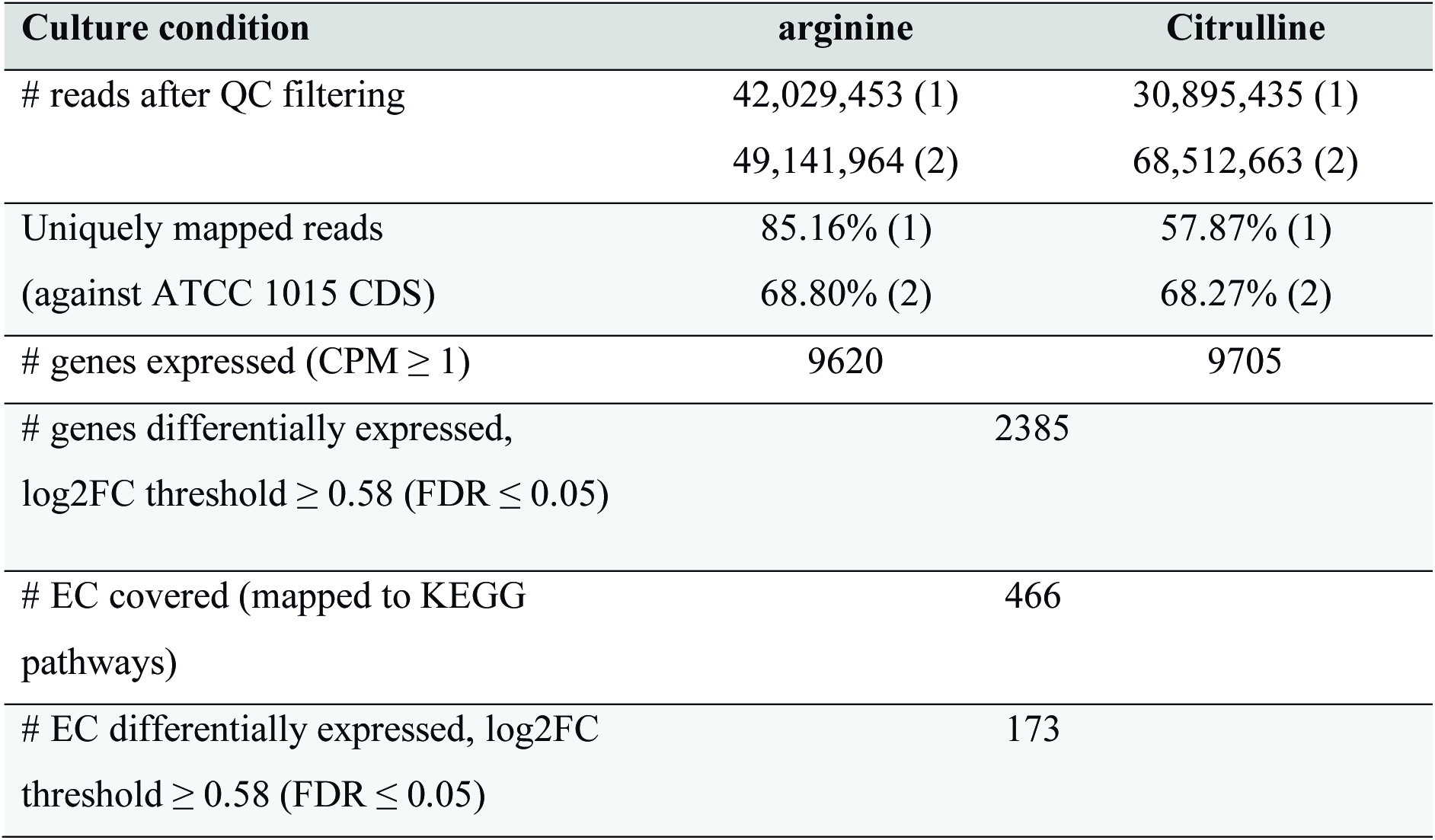
RNA seq mapping and differential expression analysis

From the 466 enzyme commission (EC) numbers that have been included in KEGG maps of metabolism [14, 15], and can be found in the annotated genome of *A. niger* ATCC 1015, 37% show differential expression. Pathway enrichment analysis shows prevalence of differentially expressed genes in pathways associated to biomass formation, such as starch and sucrose metabolism, and pathways related to amino acid biosynthesis, such as phenylalanine, tyrosine and tryptophan biosynthesis and metabolism, and valine, leucine and isoleucine biosynthesis (S3 Dataset). There is also an enrichment in differentially expressed genes related to fatty acid biosynthesis, and synthesis and degradation of ketone bodies pathways, upon addition of excess citrulline instead of arginine to the medium (S3 Dataset). Note that, although we observed higher glucose consumption in NW186 +Fe_c, glycolysis as a pathway did not show enrichment of differentially expressed genes.

Most of the pathways enriched in the arginine/citrulline supplement experiment (S3 Dataset) were also enriched in the iron experiment [7]. Phenylalanine, tyrosine and tryptophan biosynthesis, and fatty acid biosynthesis pathways showed similar behaviours in both experiments, and most of the enzymes in these pathways are down-regulated in the condition with higher citrate production.

Looking at enzymes that were connected previously to organic acid production in filamentous fungi [16], we found that alternative oxidase (AOX, or non-electrogenic ubiquinol oxidase, EC 1.10.3.11) was up-regulated in the condition corresponding to the highest citrate secretion in both experimental setups (Fig 2, S2 Dataset). It has been reported that AOX is required for efficient *A. niger* citrate production to avoid excess production of ATP [17]. However, only in the supplement experiment, up-regulation of AOX was accompanied with down-regulation of cytochrome c reductase, whereas in the iron experiment, this was not the case (Fig 2, S2 Dataset). It was found that incubating detached roots of the plant *Poa annoa* with citrate increased protein concentration of AOX without actually increasing the activity of the alternative respiration pathway itself [18]. The authors of the study hypothesised that the chelating properties of citrate might lead to the withdrawal of the Fe in the active centre of AOX, thereby rendering the protein inactive and evoking increased transcription of AOX to compensate for the inactive protein. Therefore, the transcriptional increase of *aox* observed in our two experimental setups might be due to the need to avoid excess ATP production only in the arginine/citrulline supplement experiment, whereas in the iron experiment, it might be due to high intracellular accumulation of citrate preceding its secretion in NW186 -Fe_a. Interestingly, iron limitation in itself is not enough to evoke this response, but the presence of an iron chelator seems to be necessary, *e.g*. citrate or *o*-phenanthroline (as shown for the yeast *Hansenula anomala* [19]).

**Fig 2.**
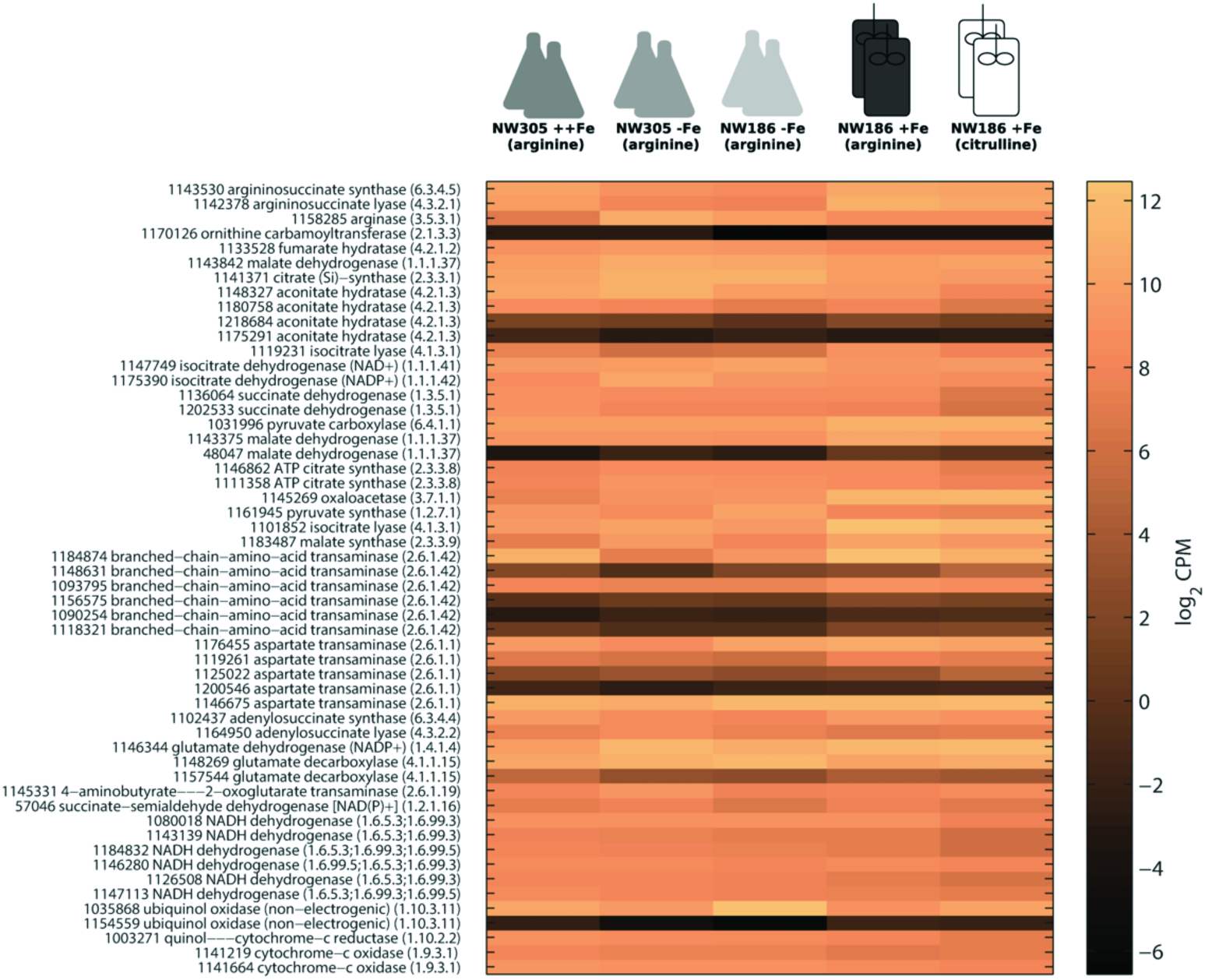
Gene expression levels (log_2_ CPM) of enzymes involved in citrate metabolism, either by direct association or as reported previously in literature. ATCC 1015 protein identifiers are given next to the enzyme names. Included are enzymes comprising the ornithine and TCA cycles, enzymes involved in amino acid turnover, and components of respiration.

Interestingly, the enzymes converting citrulline to arginine showed no differential expression in the supplement experiment (Fig 2, S3 Dataset). Citrulline, when added in excess, will not only supplement the *ΔargB* mutation, but also act as nitrogen source, thereby alleviating nitrogen limitation. Highly expressed enzymes connected to nitrogen limitation and amino acid catabolism (discussed in [16]) were down-regulated in NW186 +Fe_c compared to NW186 +Fe_a (Fig 2). In our experimental setup, increased citrate production was thus not due to nitrogen limitation in either of the two experiments; In NW305 -Fe_a and NW186 -Fe_a, iron was the limiting factor [7], whereas addition of excess citrulline in NW186 +Fe_c provides an extra source of nitrogen. However, in contrast to the general consensus, the accompanying increased biomass formation did not come at the expense of citrate yield (Table 2). The question remains why addition of excess citrulline had such a strong effect on *A. niger* citrate productivity (Fig 1B) and to what extent this is linked to changes in the amino acids pool under citrate producing conditions [20].

Analysis of the expression of enzymes involved in citrate metabolism revealed quite a contrasting transcriptomic landscape in the two experimental setups (S4 Fig). While citrate biosynthesis genes are up-regulated in the condition corresponding to increased extracellular citrate secretion in the iron experiment, there is down-regulation of citrate metabolising enzymes in the condition corresponding to increased extracellular citrate accumulation in the arginine/citrullline supplement experiment. These contrasting transcriptomic landscapes (S4 Fig, simplified in Fig 3) further support the initial hypothesis that there are two distinct mechanisms underlying the phenotype of increased citrate accumulation in the two experimental setups. Triggering increased citrate secretion by limiting Fe availability in the medium results in the highest citrate yields per biomass produced and substrate consumed (Fig 1B, Table 2). However, although efficient in terms of yield per biomass produced and substrate consumed, absolute citrate productivity (g·L^-1^·h^-1^) remains low (Fig 1B, Table 2).

**Fig 3.**
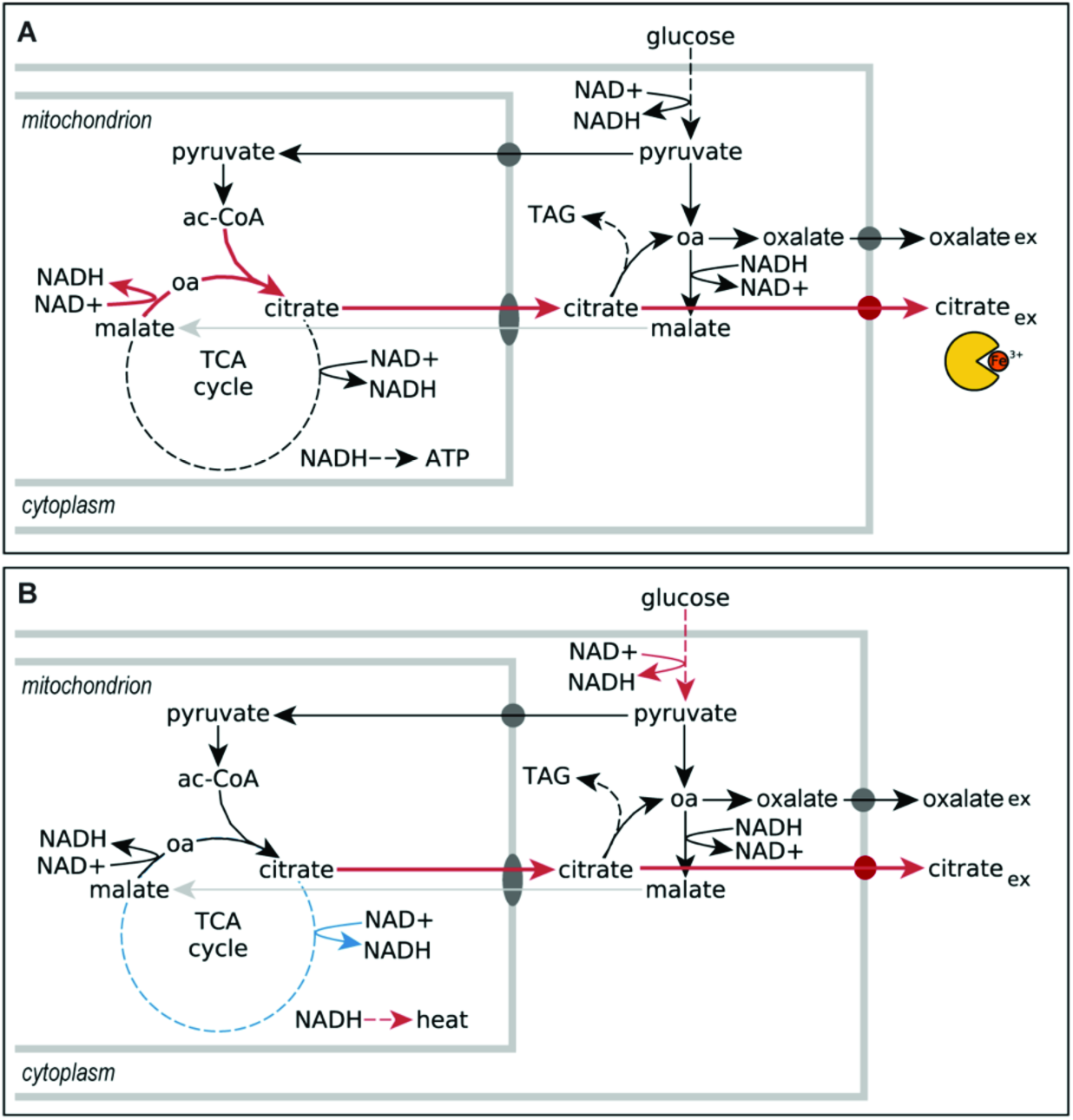
Simplification of two proposed mechanisms underlying *A. niger* citrate production. Red arrows = high flux, and blue arrows = low flux. A) Citrate as biological asset: Under iron limitation, citrate biosynthesis appears to be actively up-regulated, consistent with a possible role for citrate as *A. niger* iron siderophore [7]. B) Citrate as overflow metabolite: The fast glucose consumption in the condition where *A. niger* is supplemented with excess citrulline leads to high glycolytic flux, which in turn results in excess production of NADH. Removal of citrate from the TCA cycle prevents further production of NADH downstream of citrate. In addition, NADH re-oxidation has to be de-coupled from ATP production *via* the alternative oxidase (AOX) respiration pathway.

In the supplement experiment, we observed a higher glucose consumption rate when NW186 was supplemented with excess citrulline (Fig 1B), although this was not reflected in up-regulation of glycolysis at transcript level (S3 Dataset). A high glucose consumption rate results in high glycolytic flux that will lead to excess NADH, the turn-over of which might be limited by the capacity of further biosynthetic processes [1]. The down-regulation of cytochrome-dependent respiratory enzymes, and the up-regulation of alternative oxidase (AOX, or non-electrogenic ubiquinol oxidase) when supplementing NW186 with excess citrulline, indicate a switch from cytochrome-dependent respiration to alternative respiration mediated by AOX [1]. This effectively decouples ATP generation and NADH re-oxidation, as the reduction of molecular oxygen to water by AOX bypasses proton translocation *via* the oxidative phosphorylation complexes III and IV, resulting in a lower ATP yield [21]. Down-regulation of TCA cycle enzymes downstream of citrate in NW186 +Fe_c prevents citrate to be further metabolised, which might be a mechanism to prevent further generation of NADH [22].

### *A. niger* transporter identification and initial validation of promising candidates in yeast

To assemble a list of promising citrate transporter candidates, we combined the gene expression data obtained from the two experimental setups with three complementary computational approaches (Fig 1A). First, we looked at the problem in the context of citrate as iron siderophore. Fungal iron homeostasis genes have been found to be clustered in the genome [23], and it is not uncommon that fungal transporters are encoded by genes in close proximity to the genes encoding biosynthesis of their transport target [24]. Thus, our first selection of citrate transporter candidates were expressed genes located around *citA* in the ATCC 1015 genome, and which encoded proteins with at least one predicted transmembrane helix (tmh) domain (S5 Dataset, Sheet “cluster_approach”).

Next, we constructed two hidden Markov models (HMM) from multiple sequence alignments of biochemically characterised citrate transporters obtained from the UniProt database [25]. The sequences selected to build the models matched the terms “citrate transport” and “GO:0015137: citrate transmembrane transporter activity”, and differences in the selected sets reflect differences in protein annotation in UniProt. HMMs represent multiple sequence alignments as position-dependent scoring systems, allowing variable conservation levels and alignment length [26]. HMMs have been successfully used to identify glucose and xylose transporters in *A. niger* [27, 28]. However, construction of a HMM with reliable predictive power depends on the availability of biochemically characterised proteins with the function of interest. The HMMs were used to identify and score new citrate transporter candidates in the ATCC 1015 *in silico* proteome (S5 Dataset, Sheet “HMM_approach”). The highest scoring candidate, with protein ID 1141368, also fits with the previous approach, as it is located very close to *citA* (S5 Dataset, Sheet “cluster_approach”). Another high scoring protein, with protein ID 1155853, is also in the genomic vicinity of *citA*, although it had a much lower HMM score. However, due to biases in the sequences used to build the HMMs towards various other cell-organelles besides the plasma-membrane, and sequences of bacterial rather than eukaryotic origin, the results from the HMM approach can well be used to condense a list of likely citrate transporters across various cell-organelles, but is not suitable to pinpoint plasma-membrane *A. niger* citrate exporter candidates.

Last, we identified all plasma-membrane proteins and selected them attending to the presence and similarity of homologues in other genome sequenced *Aspergilli* or yeast that are known to either produce (*A. kawachii, Yarrowia lipolytica*) or not produce (*A. flavus, A. terreus, S. cerevisiae*) citrate (S5 Dataset, Sheet “homology_approach”). As indicated, we combined these three approaches with the expression levels and predicted localisation of the identified proteins, and finally ranked these transporters based on the expression data obtained from our two experimental setups.

Based on our pre-selection (S5 Dataset), we chose two proteins (protein IDs 1165828 and 212337) as two likely citrate exporter candidates for further validation in *S. cerevisiae*. The protein with ID 1165828 is a major facilitator superfamily transport protein with twelve transmembrane helical regions (TMHs), and is the top-ranking citrate exporter candidate based on its expression levels in the two experimental setups, and its higher similarity to transporters in *A. kawachii* and *Y. lipolytica* compared to the non-citrate producing fungi (S5 Dataset, Sheet “homology approach”). The protein with ID 212337 has 4 TMHs, and was chosen based on being the only transporter candidate in the close vicinity of *citA* that is predicted to be located in the plasma-membrane, although its low expression levels suggest lesser importance as actual citrate exporter (S5 Dataset, Sheet “cluster_approach”).

*S. cerevisiae* was transformed with plasmids containing either one of the putative citrate exporter candidate genes under the control of a copper inducible promoter (CUP1). In the first growth experiment, we induced expression of the transporter proteins after 4 h with 1 mM CuSO_4_, and found that neither the untransformed control strain, nor the strain transformed with the protein 212337 encoding gene, secreted any citrate. However, the strain transformed with the protein 1165828 encoding gene, hereafter named *citT*, accumulated a small amount of extracellular citrate, indicating that this could be a citrate exporter. To further verify our initial results, we grew *S. cerevisiae* transformed with the *citT* gene with either glucose or glycerol in the medium, and either did or did not induce expression with CuSO_4_. Reconfirming our initial observations, we found measurable accumulation of citrate in the extracellular medium only in *S. cerevisiae* in which *CitT* expression was induced, but not in the non-induced transformant, nor in the untransformed parent strain (Fig 4).

**Fig 4.**
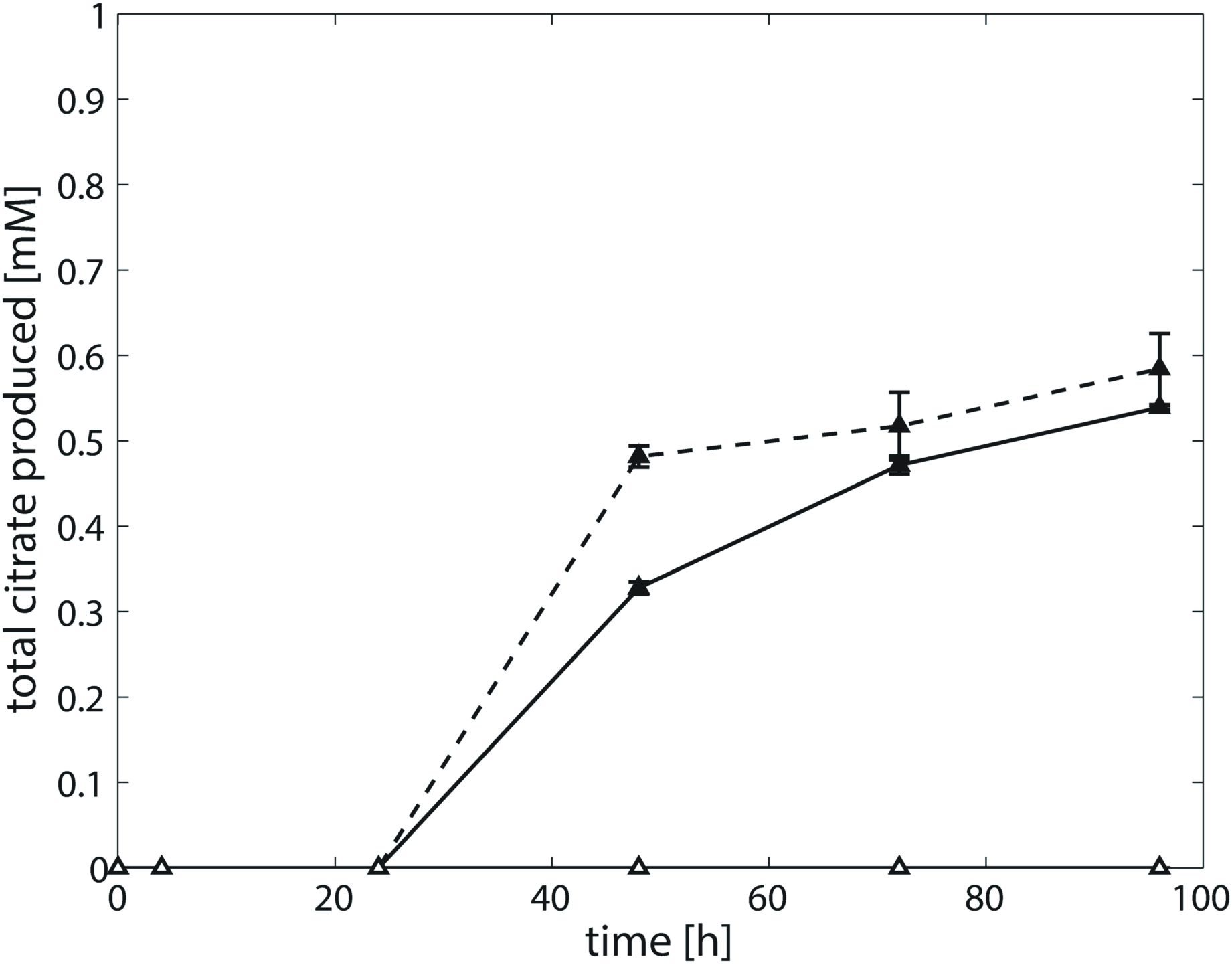
Citrate secretion in *S. cerevisiae* transformed with CitT. Solid line = growth on glucose, dashed line = grown on glycerol, filled symbols = induced, empty symbols = non-induced. Note that there was no measurable extracellular citrate accumulation in the untransformed parent strain. Measurements indicate the average of two biological replicates.

### Deletion of *citT* abolishes citrate production in *A. niger*

Accumulation of citrate in the medium of *S. cerevisiae* expressing *citT* strongly suggests that the encoded protein can function as a citrate exporter. To understand the molecular consequences of a *citT* deletion, the endogenous *A. niger* gene was replaced by the selection marker *pyrG* from *A. oryzae*. Three independent *ΔcitT* strains, labelled *ΔcitT-1.1, ΔcitT-2.1* and *ΔcitT-4.1*, were selected for further study.

The phenotypes of the *ΔcitT* strains were observed on MM plates supplemented with glucose as carbon source. In comparison to the wildtype N402, there were no differences of phenotypes and growth rates. Moreover, the *ΔcitT* strains were able to grow normally on medium containing citrate as sole carbon source. That means the deletion of the *citT* gene is not lethal to *A. niger*, and that citrate import is not affected by the *citT* deletion.

Shake flasks were used to screen the *ΔcitT* strains for their ability to secrete citrate. Glucose consumption, organic acid concentrations and pH were followed in time (Fig 5). Using a starting pH of 3.63, all strains acidified the medium. In case of the N402 control strain, the pH decreased to 1.72 at 48 h and then increased slightly to pH 1.83 at 96 h (Fig 5A). The pH of the *ΔcitT* strains were constant at pH 1.70 from 48 h onwards (Fig 5B-D). Glucose consumption of the *ΔcitT* strains was slower than that of the wildtype; in the supernatant of the *ΔcitT* strains, glucose was depleted only after 72 h, whereas glucose was already depleted after 48 h in the supernatant of N402. Most of the glucose was converted to oxalate in the *ΔcitT* strains, and only the wildtype strain exported citrate. The highest amount of citrate detected was ~1.3 g·L-^1^ (at t = 48 h). After 24 h, the oxalate yields in the *ΔcitT* strains were around 10% higher than those of the N402 strain. These results reconfirm that the *citT* gene encodes an *A. niger* citrate exporter. The three *ΔcitT* strains showed the same results, so only *ΔcitT-1.1* was selected for further studies.

**Fig 5.**
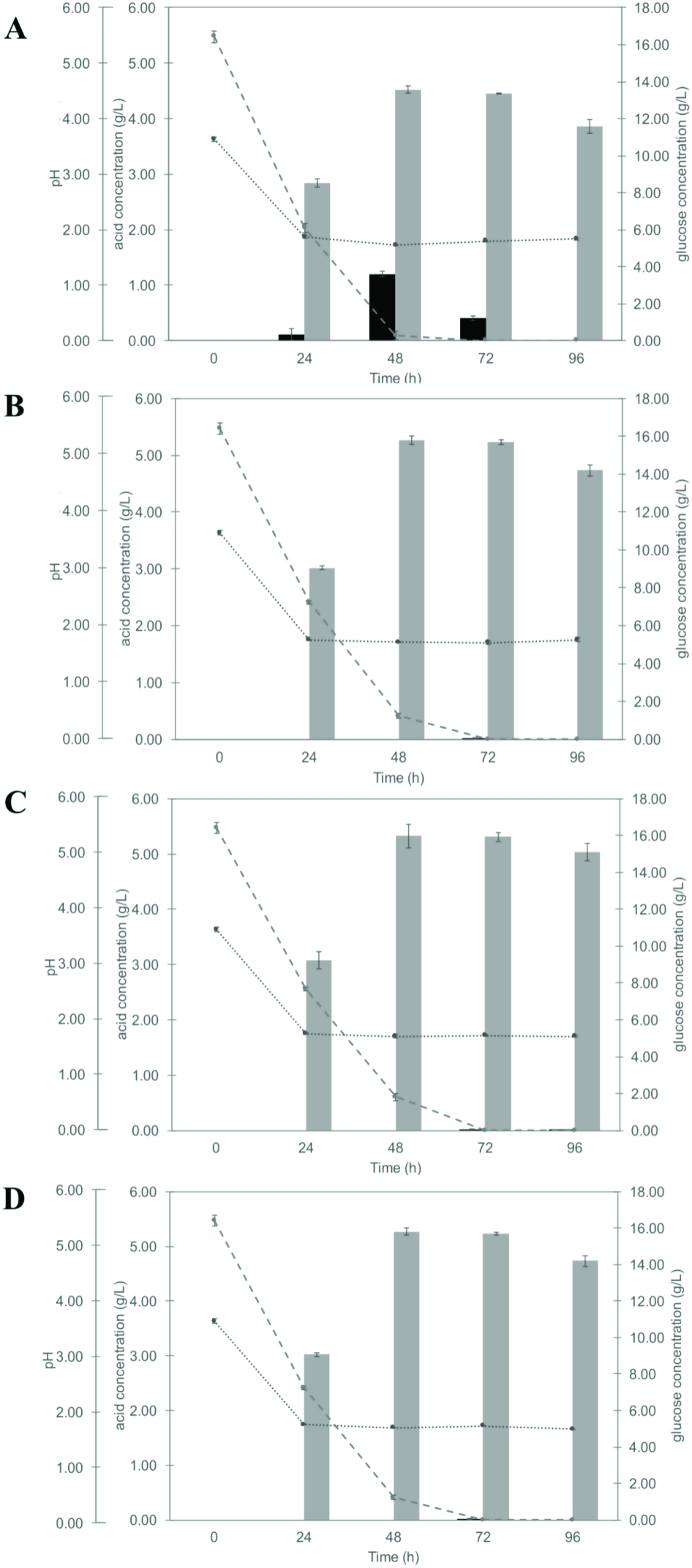
Screening citrate exporter knock-out strains by small-scale fermentation using glucose as carbon source. The graphs show glucose consumption, organic acid production, and pH of *A. niger* strain A) N402, B) *ΔcitT-1.1*, C) *ΔcitT-2.1* and D) *ΔcitT-4.1*. Black bar = citrate, grey bar = oxalate, dotted line = pH, and dashed line = glucose. Measurements indicate the average of two (*ΔcitT strains)* or three (N402) technical replicates.

As initial screen, the shake flask experiments were not optimised for citrate production. To further understand the consequences of a *citT* deletion for citrate production, we used a medium with 120 g·L^-1^ glucose [29]. For citrate production, a pH below 3.0 is crucial [6, 30]. The pH of the fermentation was maintained at 2.5. At this pH, we can still monitor the amount of oxalic acid produced as it high enough to prevent extracellular conversion of oxalic acid to formic acid by oxalate decarboxylase [7, 31] and low enough to inactivate extracellular gluconic acid formation.

In these experiments, extracellular citrate accumulation was detected only in the supernatants of the wildtype, where the 6.7 fold increase in glucose drastically increased the citrate productivity of the N402 strain to 30 g·L^-1^ (versus 0.10±0.04 g·L^-1^ for the *ΔcitT* strain (Fig 6)). It should be noted that hyper producing strains, reaching over 100 g·L^-1^, have been evolved from very similar genetic background. The resulting strains have structural genomic re-arrangements but with the same set of genes, except for mutations in genes involved in citrate production [32].

**Fig 6.**
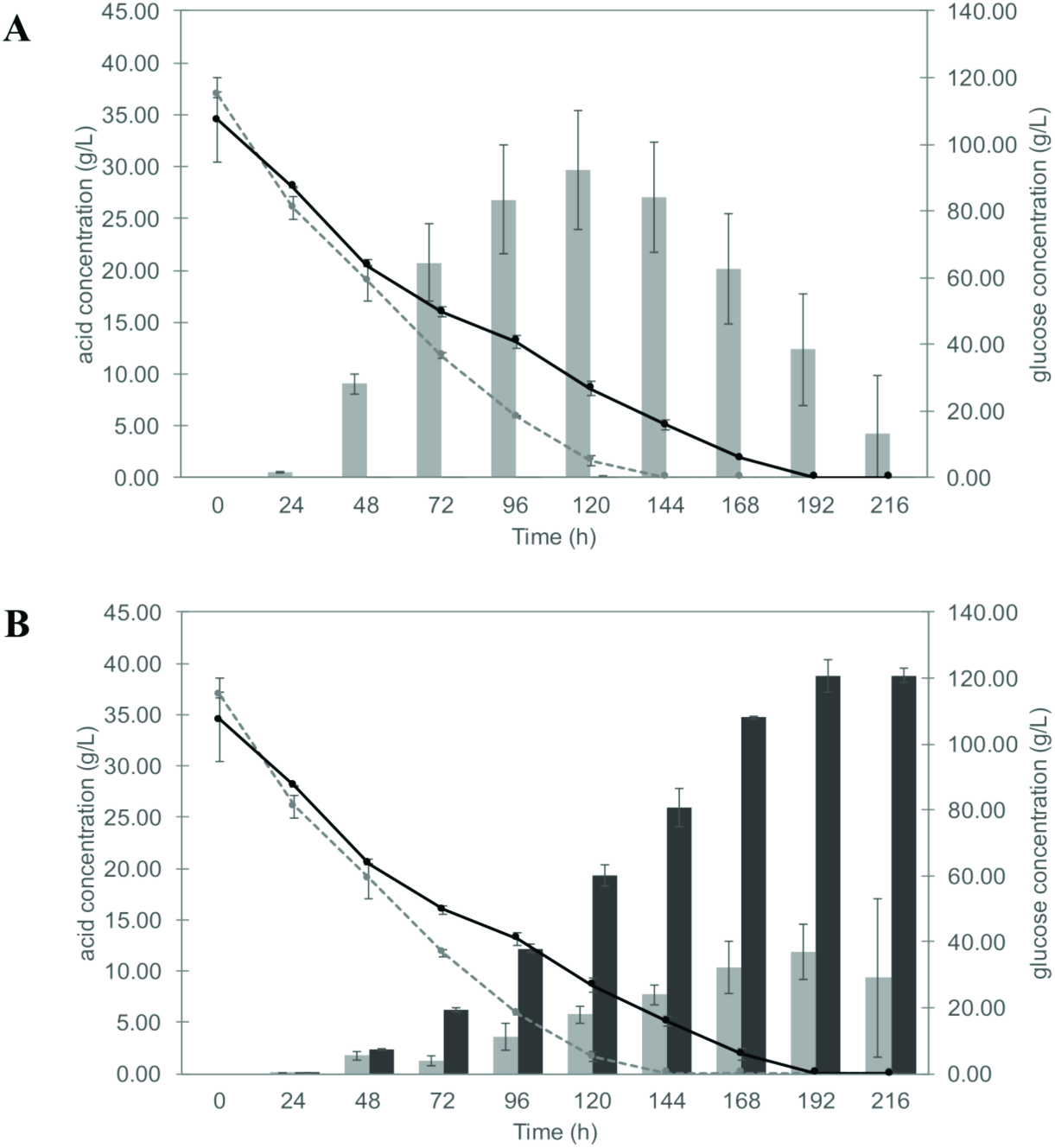
Fermentation of *A. niger* N402 and the *ΔcitT* strain under conditions more optimal for citrate production. The concentration of A) citrate and B) oxalate are represented as bar graphs. Grey bar = N402, black bar = *ΔcitT*, dotted line = glucose in N402 supernatant, and bold line = glucose in *ΔcitT* supernatant. Measurements indicate the average of three replicates.

Another observation was that the *ΔcitT* strain took 192 h to deplete glucose, while the wildtype strain took only 144 h (Fig 6). *A. niger* produces citrate *via* glycolysis and the subsequent mitochondrial TCA cycle. In the cytosolic glycolysis, glucose is converted to pyruvate, which is subsequently transferred to the mitochondrion. In the mitochondrion, pyruvate is converted to acetyl-CoA and then acetyl-CoA and oxaloacetate are condensed to citrate, which is transported to the cytoplasm. At this point, citrate can either be converted to oxaloacetate by ATP-citrate lyase, which can then be directly converted in oxalate by oxaloacetate acetylhydrolase [33], or secreted *via* the citrate exporter (Fig 3). Upon deletion of the citrate exporter gene, the *ΔcitT* strain can no longer secrete citrate. In these strains, glucose consumption is reduced, suggesting that the high glycolytic flux observed under citrate producing conditions and induced by high initial glucose concentrations in the medium cannot be maintained. This might be due to an inhibitory effect of an increased intracellular citrate pool on phosphofructokinase [34]. Deprived of the possibility to export citrate, the *ΔcitT* strain secreted other organic acids. Oxalate was found in the supernatants of the *ΔcitT* strain with the maximum amount around 40 g·L^-1^, while the wildtype produced 12 g·L^-1^ at 192 h under these conditions (Fig 6B).

## Conclusion

In the present work, we compared RNA expression levels of two experimental setups in which a relative increase in *A. niger* citrate secretion was triggered by either limiting iron availability or supplementing the fungus with excess citrulline instead of arginine. Our analyses of the underlying transcriptomic landscapes show that extracellular citrate accumulation can be the answer to multiple environmental conditions. As such, we propose that there is no such thing as *the* mechanism underlying *A. niger* citrate production and secretion. In combining the expression data of both the iron and supplement experiments with three different *in silico* approaches, we condensed the *A. niger* transportome into a shortlist of the most promising citrate transporter candidates. The candidate genes were tested in *S. cerevisiae*. While one of the two candidates produced no significant change on the production phenotype, the other one enabled citrate secretion in this organism, thus leading to the identification of an *A. niger* citrate transporter, CitT. *A. niger* knock-out strains of the corresponding citrate exporter gene secrete negligible amounts of citrate, even under conditions optimised for comparatively high citrate production. Although we cannot rule out that another transporter is responsible for citrate secretion under industrial hyper-producing conditions, our results suggest a single major facilitator superfamily transport protein is responsible for active *A. niger* citrate export.

## Materials and methods

### Strains and Media

*Escherichia coli* DH5α was used for standard gene cloning purposes. *E*. *coli* was cultured in Luria broth (LB) medium (5 g·L^−1^ yeast extract, 10 g·L^−1^ peptone, 10 g·L^−1^ NaCl) supplemented 100 mg·L^−1^ ampicillin when necessary.

*S. cerevisiae* strain CENPK2-1D (MATα; his3D1; leu2-3_112; ura3-52; trp1-289; MAL2-8c; SUC2) was used for testing the citrate transporter candidates. Preparation of CENPK2-1D yeast electro-competent cells and transformation was performed according to Suga and Hatakeyama [35]. Yeast transformed cells were selected in synthetic media (SD) with 2% (w/v) dextrose, 0.67% (w/v) Yeast Nitrogen Base without amino acids (BD), 0.14% (w/v) Yeast Synthetic Drop-out Medium supplement without uracil, tryptophan, histidine and leucine (Sigma-Aldrich), 0.0076% (w/v) histidine, 0.0076% (w/v) tryptophan and 0.038% (w/v) leucine. For *S. cerevisiae* transformation, the cell suspension was mixed with plasmid or linear DNA, transferred to a pre-chilled cuvette (0.2 cm Gene Pulse, Bio-Rad), pulsed at 2.5 kV, 25 μF, 200 Ω using Gene Pulser Xcell (Bio-Rad) and plated on SD agar-plates.

*A. niger* strain NW305 (*cspA, goxC17*, Δ*argB*) [36], and NW186 (*cspA1, goxC17, prtF28, ΔargB*) - a derivative of NW185 [12] with restored uridine prototrophy [7] - were used to study conditional citrate export. *A. niger* strain N402 was used as wildtype strain. *A. niger* strain MA169.4 (*kusA*::DR-*amdS*-DR, *pyrG^-^*) was used as recipient strain in the knock-out transformation experiment [37] using the *pyrG* gene from *A. oryzae* as a selectable marker gene. The pWay-pyrA plasmid was used as a control for transformation. Strains were maintained on complete medium (CM) containing 1 g·L^-1^ casamino acid, 5 g·L^-1^ yeast extract, 20 mL·L^-1^ ASPA + N [38], 1 mL·L^-1^ Vishniac solution [39], 1 mM MgSO_4_ and supplemented with 5 g·L^-1^ glucose as carbon source, at 30°C [38] and supplemented with 0.2 g·L^-1^ uridine and/or 0.02% arginine when necessary. *A. niger* spores were harvested with Saline-Tween (0.9% NaCl and 0.001% Tween80).

### Heterologous expression of putative transporter genes in *S. cerevisiae*

Primers used for cloning are listed in S6 Table. The coding sequences of two putative citrate transporter genes, with ATCC 1015 protein IDs 1165828 and 212337, were amplified from *A. niger* N402 genomic DNA. The amplified fragments were introduced into a derivative of the pYES-plasmid, whereby the GAL1 inducible promoter was replaced by the CUP1 inducible promoter [40]. The plasmid was PCR-amplified with the Q5 polymerase following the manufacturer’s protocol. The coding sequence of the citrate exporter candidate 1165828 was cloned using Golden Gate. The Golden Gate reaction mixture was prepared as follows: 400 U of T4 DNA ligase (NEB), 10 U of BsmBI (NEB), 1.5 μL of BSA (1 mg·mL^-1^), 1.5 μL of T4 DNA ligase buffer (NEB), ~40 fmol of each PCR product, and water to bring the volume up to 15 μL. The reaction mixture was incubated in a thermocycler according to the following program: 37°C for 10 min prior to 25 cycles of digestion-ligation (37°C for 3 min, 16°C for 4 min) followed by a final digestion step (55°C for 10 min) and a heat inactivation step (80°C for 10 min). *E. coli* DH5α competent cells were transformed with 1 μL of the digestion-ligation reaction, and transformants were selected on ampicillin plates. Plasmid extraction was performed using the GeneJET Plasmid Miniprep kit (Thermo Fisher Scientific) following the manufacturer’s instructions. CENPK2-1D competent cells were transformed with a sequence-verified plasmid. The coding sequence of the citrate exporter candidate 212337 was cloned using the yeast homologous recombination assembly strategy. For the in-yeast assembly, CENPK2-1D cells were directly transformed with 100 ng of the PCR-amplified pYES-CUP1 plasmid and an equimolar amount of insert, and plated on selective medium. Positive transformants were picked and grown overnight in liquid medium. Plasmids were extracted using the GeneJET Plasmid Miniprep kit following the manufacturer’s instructions with minor modifications. Lysis was performed by transferring the cells to Lysing MatrixC 2 mL tubes (MP Biomedicals) and homogenising them for 40 s with a FastPrep-24 from MP Biomedicals. *E. coli* DH5α competent cells were transformed with the extracted plasmids for propagation, and the plasmids were then again extracted using the GeneJET Plasmid Miniprep kit following the manufacturer’s instructions, and sequenced for insert verification.

For the growth experiments, yeast transformant strains were pre-grown in 15 mL SD medium supplemented with 0.0076% uracil to allow growth of the parent strain. 300 μL of the pre-growth culture were inoculated in 100 mL shake flasks containing 20 mL SD medium described above, containing either 20 g·L^-1^ glucose or 20.44 g·L^-1^ glycerol. For CUP1 promoter-induction, CuSO_4_ (final concentration 1 mM) was added after 4 h of growth.

### *A. niger* transporter identification and validation

#### Supplement experiment for transporter identification

For pre-growth of *A. niger* NW186 and NW305, a total of 1ˣ10^6^ spores·mL^-1^ were inoculated in 1L Erlenmeyer shake flasks containing 200 mL medium with the following composition:1.2 g·L^-1^ NaNO_3_ or 0.93 g·L^-1^ (NH_4_)_2_SO_4_, 0.5 g·L^-1^ KH_2_PO_4_, 0.2 g·L^-1^ MgSO_4_**·**7H_2_O, 40 μL·L-^1^ Vishniac solution, and supplemented with 50 g·L^-1^ (~250 mM) glucose as carbon source, and either 1.1 mM (0.2 g·L^-1^) arginine or 5 mM (0.88 g·L^-1^) citrulline. After 24 h of pre-growth, 11 g of *A. niger* mycelium was transferred to controlled fermentors, containing the same medium as for the pre-growth. The supernatants were collected for determination of extracellular metabolite concentrations, especially carbon sources and organic acids, by high-performance liquid chromatography (HPLC) as described previously [7].

#### Construction of knock-out strains for transporter validation

*A. niger* citrate exporter knock-out strains were constructed using the split-marker approach [41]. The c*itT* gene was deleted from the MA169.4 genome (isogenic with N402), which is defective in the Non-Homologous End-Joining (NHEJ) pathway through a transiently silenced k*usA g*ene [42]. The experimental steps representing the construction of the knock-out strains can be found in S7 Fig. Correct deletion of the gene was confirmed with PCR. Primers used in this experiment are in S6 Table.

A protoplast-mediated transformation of *A. niger* MA169.4 was performed by adapting the protocol of [38]. A total of 1ˣ10^6^ spores·mL^-1^ was inoculated in a 1L Erlenmeyer flask coated with 5% dimethydichlorosilane in heptane, containing 250 mL CM supplemented with 10 mM uridine. The culture was incubated at 30°C and 100 rpm for 12 h. The mycelium was harvested by filtration through sterile miracloth and washed once with SMC buffer. Lysing enzymes from *Trichoderma harzianum* (Sigma) were used for protoplast formation (400 mg enzymes per g mycelium). Mycelium was incubated at 37 °C and 80 rpm for 3 h. Protoplasts and mycelium were separated by filtration through the miracloth. The filtrate containing the protoplasts was centrifuged at 2000 g and 10 °C for 10 min, washed twice with 1 mL STC buffer, re-suspended in 1 mL STC buffer, and aliquoted to 100 μL per reaction. After transformation, single *A. niger* transformant colonies were purified, and the transiently silenced *kusA* gene was restored on MM plates containing fluoroacetamide (FAA) as described in [37].

#### Initial screening of ΔcitT candidate knock-out strains for citrate secretion

For initial screening, shake flasks were used. The wild type N402 and the *ΔcitT* strains were pre-cultured in CM containing 1ˣ10^6^ spores·mL^-1^ at 30 °C and 200 rpm. After 18 h, mycelium was harvested and then rinsed with equal amounts of water. Mycelium (5 g wet weight) was transferred to 200 mL medium supplemented with 18 g·L^-1^ glucose or 10.5 g·L^-1^ citrate as carbon sources (medium adjusted to pH 3.5), and then incubated at 30 °C, 200 rpm.supernatant was collected every 24 h to determine the amount of carbon source consumption and organic acid production by HPLC.

#### *Citrate production of* Δ*citT:*

Growth experiments for determination of Δ*citT c*itrate production were performed in fermentors in triplicate. Pre-cultures of the N402 and a Δ*citT* strain were prepared with 1ˣ10^6^ spores·mL^-1^ inoculated in 200 mL CM-SLZ, containing 1 g·L^-1^ casamino acid, 5 g·L^-1^ yeast extract, 3 g·L^-1^ (NH_4_)_2_SO_4_, 1 g·L^-1^ K_2_HPO_4_, 1 g·L^-1^ KH_2_PO_4_, 0.5 g·L^-1^ MgSO_4_·7H_2_O, 0.014 mg·L^-1^ MnSO_4_·5H_2_O, 0.01 mg·L^-1^ FeCl_2_·6H_2_O, 0.075 μg·L^-1^, ZnCl_2_, 1 mL·L^-1^ and 10 g·L^-1^ glucose, adjusted to pH 5.5, and incubated at 30 °C, 200 rpm, for 18 h. Mycelium was harvested and then rinsed with equal amounts of water and 10 g (wet weight ~ 0.92 g CDW) of mycelium was transferred to 750 mL of SLZ medium, containing 3 g·L^-1^ (NH_4_)_2_SO_4_, 1 g·L^-1^ K_2_HPO_4_, 1 g·L^-1^, KH_2_PO_4_, 0.5 g·L^-1^ MgSO_4_·7H_2_O, 0.014 mg·L^-1^ MnSO_4_·5H_2_O, 0.01 mg·L^-1^ FeCl_2_·6H_2_O, 0.075 μg·L^-1^ ZnCl_2_, and 120 g·L^-1^ glucose, and adjusted to pH 2.5 [29]. Methanol (2% (v/v)) was added as a promotive element for citrate production [43] and 30% (v/v) polypropylene glycol (ppg) was added as an anti-foaming agent. The cultures were aerated with sterile compressed air continuously blown through the sparger at a rate of 0.6 slpm and the stirring speed was adjusted at 1000 rpm. The pH was maintained at 2.5 by 5 M NaOH, and the temperature was set at 30 °C.

### Bioinformatic analysis for transporter identification

#### RNA isolation, sequencing and RNAseq data processing and analysis

Total RNA extraction, RNA sequencing and RNAseq data processing and statistical analysis of the supplement experiment were performed as described in [7]. The aligned .bam files were submitted to the European Nucleotide Archive (ENA) under the accession number PRJEB24704. Data was compared to data from Odoni *et al* [7] (iron experiment), submitted to the European Nucleotide Archive (ENA) under the accession number PRJEB20746.

#### Shortlisting putative citrate transporter candidates

To assemble a list of likely citrate transporter candidates, three complementary approaches were followed. RNAseq analysis was supplemented with (i) a cluster approach, (ii) a HMM approach, and (iii) a homology based approach. In (i), we analysed protein coding genes in close proximity to *citA* for features that make them putative transporter protein candidates; proteins with transmembrane helix structures (from the *A. niger* ATCC 1015 *in silico* proteome) were identified using the stand-alone TMHMM 2.0 software package [44, 45]. Protein localisation was predicted using the stand-alone protComp software (www.softberry.com). (ii) Hidden Markov models were built as described [27] from two sets of proteins (“citrate transport” and “GO:0015137”) downloaded from the UniProt database [25]. (iii) The genomes used for the homology approach were downloaded from the JGI database [46]: *Aspergillus kawachii [47], Aspergillus nidulans* [48, 49], *Aspergillus flavus [48], Aspergillus fumigatus [50-53], Aspergillus terreus [48], Yarrowia lipolytica [54]* and *Saccharomyces cerevisiae* [54].

## Supporting information captions

**S1 Dataset.** Expression of all CDS

**S2 Dataset.** DE suppl Fe CDS

**S3 Dataset.** Pathway Maps ratio

**S4 Fig.** Differential expression of genes encoding enzymes involved in *A. niger* citrate metabolism. Enzymes and pathways are colour coded according to their transcriptional changes in A) the iron experiment, and B) the supplement experiment. Red colour indicates higher expression in the condition with higher citrate production (-Fe in the iron experiment, citrulline in the supplement experiment) compared to the condition that had a lower citrate per glucose production (++Fe in the iron experiment), or total citrate production (arginine in the supplement experiment). Similarly, blue indicates lower expression in the conditions with either a higher citrate per glucose yield, or higher total citrate production. Grey indicates no significant (FDR ≤ 0.05 and log2FC ≥ 0.58) changes in expression. Abbreviations: SB = siderophore biosynthesis, oa = oxaloacetate.

**S5 Dataset.** Citrate transporter candidates

**S6 Table.** Primers used in this study.

**S7 Fig.** Scheme representing experimental steps for deletion of *citT* gene from the *A. niger* genome.

